# Deterministic and probabilistic fate decisions co-exist in a single retinal lineage

**DOI:** 10.1101/2022.08.11.503564

**Authors:** Elisa Nerli, Jenny Kretzschmar, Tommaso Bianucci, Mauricio Rocha-Martins, Christoph Zechner, Caren Norden

**Affiliations:** Instituto Gulbenkian de Ciência, Rua da Quinta Grande 6, 2780-156 Oeiras, Portugal; Max Planck Institute of Molecular Cell Biology and Genetics, Pfotenhauerstraße 108, 01307 Dresden, Germany; Max Planck Center for Systems Biology, Pfotenhauerstraße 108, 01307 Dresden, Germany; Cluster of Excellence Physics of Life, TU Dresden, 01062 Dresden, Germany

**Keywords:** Neurogenesis, retina, fate decisions, competence, plasticity, zebrafish, live-imaging, modelling

## Abstract

Correct nervous system development depends on the timely differentiation of progenitor cells into neurons. While the output of progenitor differentiation is well investigated at the population and clonal level, the possibilities and constraints for fate decisions of specific progenitors over development are less explored. Particularly little is known about their variability and competence plasticity. To fill this gap, we here use long-term live imaging to follow the outcome of progenitor divisions in the zebrafish retina.

We find that neurogenic Atoh7 expressing progenitors produce different neuronal types over development with time-dependent probabilities. Interestingly, deterministic and probabilistic fate decisions co-exist in the same lineage. While interference with the deterministic fate affects lineage progression, interference with fate probabilities of the stochastic lineage branch results in a broader range of fate possibilities than seen in controls. When tissue development is challenged, Atoh7 expressing progenitors can produce any neuronal type, arguing against the concept of fixed competence windows. Stochastic modelling of fate probabilities in challenged conditions revealed a simple gene regulatory network able to recapitulate the observed competence changes during development. Based on our results, we postulate that fate plasticity could be involved in robust retinal development, a concept possibly applicable to other tissues.

## Introduction

To generate organs in a developing embryo, cells progressively become more specialized as they differentiate. Cell differentiation needs to be tightly regulated to produce the correct cell types at the right developmental time. Impairment of the temporal sequence of differentiation can have detrimental consequences for organismal development, including incorrect organ size or cellular arrangements^1, 2^. It is thus important to unveil the factors that ensure the production of the right cell type with the correct timing. This is particularly true for the formation of the central nervous system (CNS), where timely emergence of the different neurons is an important step that later ensures correct neuronal connectivity and the formation of functioning neuronal networks^2–4^. Unsurprisingly, changes in the timing of neuronal specification and differentiation can impair brain formation. This in turn can lead to severe cognitive deficits including mental retardation and impairments in motor coordination^5^. Nevertheless, particularly in vertebrates, the factors involved in different neuronal fate decisions during development are not yet fully disclosed. This is different for the Drosophila nervous system, where the temporal regulation of fate decisions has been more thoroughly explored for example in the CNS and the optic lobe^6–9^. In these areas, a defined sequence of progenitor divisions leads to the formation of the different neurons in a consecutive manner. This sequence arises during development as multipotent progenitors are competent to form different neuronal types through the sequential expression of defined transcription factors^6, 10, 11, 12–14^. In the vertebrate nervous system, however, it is less understood whether and how single progenitors competence changes during development to give rise to different neuronal types at the right time and in the right proportions. So far, some progress has been made to understand neuronal birth orders in areas including the neocortex, spinal cord, olfactory bulb and the retina^15–20^ but these studies have been mostly performed at the clonal or the population level. This eventually led to different interpretations on whether progenitors competence is pre-determined in progenitor cells, resulting in fixed competence windows during development^19, 21, 22^, or whether it is variable and influenced by stochastic processes acting on fate decision mechanisms^23–26^.

To better understand the variability and stereotypicity of fate decisions in the vertebrate CNS, a quantitative appreciation of whether and how far the fate outcome of defined progenitor divisions changes over development is needed. To date, this has been challenging due to the fact that many parts of the vertebrate brain are not easily accessible for experimental manipulation and due to the plethora of different progenitors and neuronal types inhabiting different brain areas. An attractive system to circumvent these issues is the developing retina, the part of the CNS responsible for light collection and transmission. It is located on the outside of the embryo and is populated by only five major neuronal types that are clearly distinguishable by their final position, morphology, and mode of migration^27, 28^ (Figure 1 B, B’). In addition, the imaging potential of the zebrafish embryo allows to follow the exact sequence and outcome of progenitor divisions over time and the lineage relationships between cell types in a quantitative manner^23, 29–31^. Thus, the zebrafish retina is an ideal model to elucidate the temporal dynamics of progenitor competence and consequently to assess stereotypicity and variability in lineage decisions.

**Figure 1.**
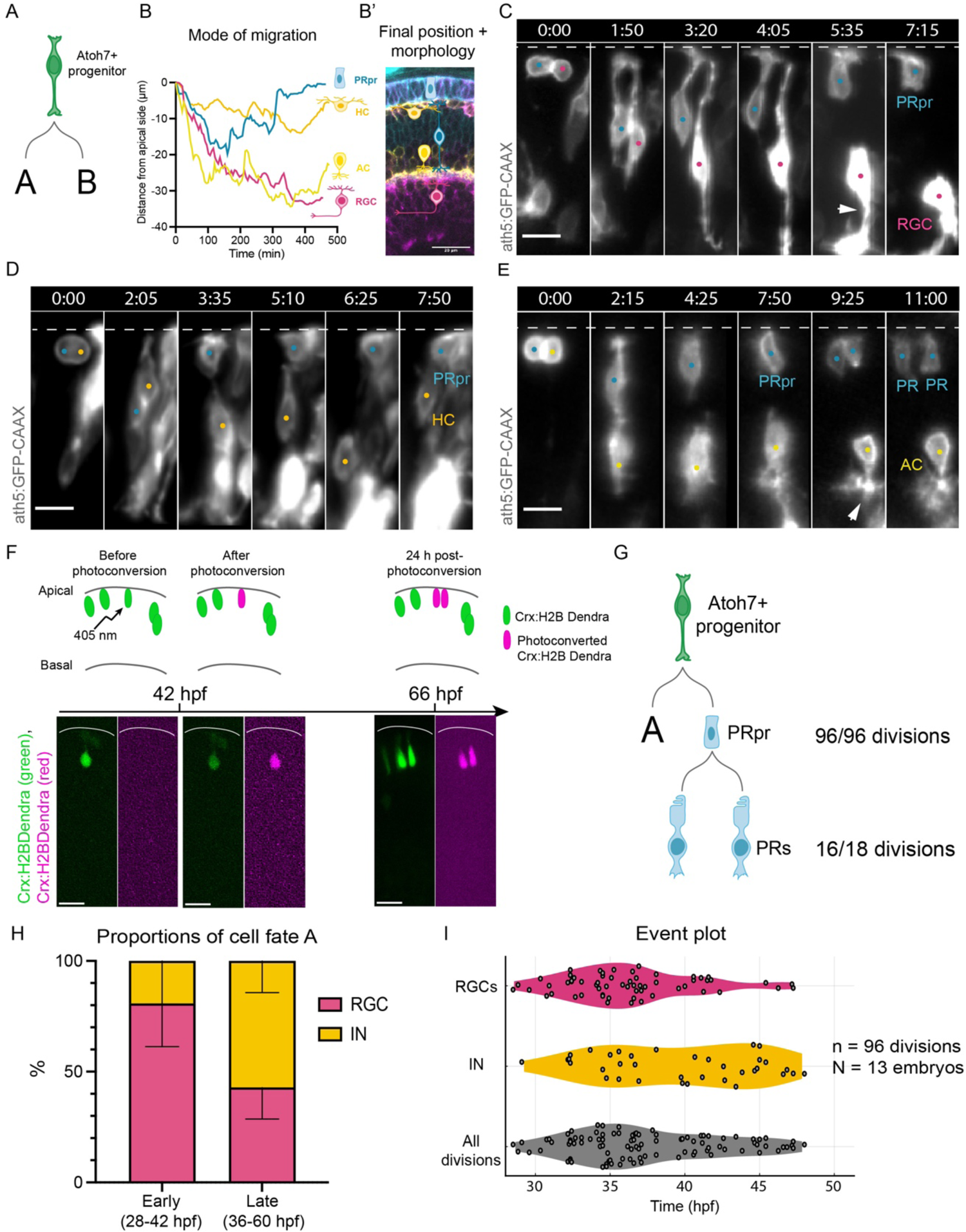
Atoh7+ progenitors divide asymmetrically to produce a PRpr and a sister cell. A) Schematic of Atoh7+ divisions. The focus of this study is to find out what daughter cells are generated in branch A and B and the spectrum of fate decision possibilities of Atoh7+ progenitors. B) Cell fate assignment strategy based on mode of migration. The plot indicates typical movement trajectories for each cell type. B’) Cell fate assignment strategy based on morphology and final neuronal position in the laminated retina. C) Montage of Atoh7+ progenitor division generating an RGC (magenta dot) and a PRpr (cyan dot). Dashed line indicates the apical side, arrowhead points to RGC axon. ath5:GFP-CAAX (Atoh7, grey), scale bar 10 µm. D) Montage of Atoh7+ progenitor division generating an HC (orange dot) and a PRpr (cyan dot). Dashed line indicates the apical side. ath5:GFP-CAAX (Atoh7, grey), scale bar 10 µm. E) Montage of Atoh7+ progenitor division generating an AC (yellow dot) and a PRpr (cyan dot). Dashed line indicates the apical side, arrowhead points to basal dendrites. ath5:GFP-CAAX (Atoh7, grey), scale bar 10 µm. F) Photoconversion experiment. A 405 laser was used to photoconvert isolated cells labelled with Crx:H2BDendra at 42 hpf. 24 hours after photoconversion, photoconverted PRs were assessed. (Top) Schematic of the experiment, (bottom) representative images of the different steps of the experiment. Crx:H2BDendra (green), photoconverted Crx:H2BDendra (magenta). Scale bar 10 µm, white line indicates apical side of the retina. G) Schematics of outcome of Atoh7+ divisions and PRpr divisions. H) Distribution of fates for cell A acquired during early and late neurogenic windows. N = 13 embryos, n = 96 Atoh7+ divisions. Mean and 95% CI are indicated. I) Event plot of all divisions analysed in (H).

We here explore the possibilities as well as the constraints of fate decisions that specific zebrafish retinal progenitors present over time. Interestingly, we find that Atoh7-expressing neurogenic progenitors consistently give rise to a photoreceptor precursor and a sister cell of different fate. While photoreceptor production is deterministic, the fate of the sister cell is acquired in a probabilistic manner and these probabilities change over development. If the deterministic photoreceptor fate decision is impaired, Atoh7 progenitors do not generate a different known neuronal fate, resulting in lineage and tissue impairments. In contrast, interference with the probabilistic branch of the lineage resulted in changes in the probabilities of generating different fates, and maintenance of structural tissue integrity. Stochastic modelling of these changed probabilities revealed a simple gene regulatory network (GRN) that faithfully predicted the proportions and timing of fate decisions during normal retinal neurogenesis.

## Results

### Neurogenic Atoh7+ progenitors produce one photoreceptor precursor and a sister cell of variable fate

Progenitors expressing the proneural transcription factor Atoh7 (*atonal bHLH transcription factor 7*, also called Ath5) give rise to most retinal neurons^30, 32–34^. However, the temporal sequence and distribution of fate decisions resulting from these divisions has only begun to be understood^30, 35^ (Figure 1 A). To quantitatively analyse fate distributions, we used an Atoh7-driven reporter construct to label these progenitors (ath5:GFP-CAAX)^36, 37^. We followed their division modes and fate outcomes by long term light sheet imaging^38^ performed between the onset of neurogenesis at 28 hours post fertilization (hpf) and 60 hpf, the time when most progenitors have entered neurogenesis^39–41^. Sister cell fate could unambiguously be assigned due to the distinct migration modes of the different emerging neurons, their morphology during and after migration and their final position within the tissue (Figure 1 B, B’, see Methods for details).

Analysis of 96 divisions from 13 embryos during the entire neurogenic window revealed that Atoh7+ progenitors divide asymmetrically and reproducibly produce one cell with columnar morphology and a sister cell of variable fate (Figure 1 C-E, Video 1).

Apically positioned cells with columnar morphology were previously suggested to be immature photoreceptors, so-called photoreceptor cell precursors (PRpr)^30, 34, 38^. These cells were characterized by the expression of *crx* (cone-rod homeobox)^42, 43^ and assumed to divide symmetrically to produce two PRs^42, 43^. However, whether all PRs result from such committed precursor divisions remained unclear. We found that the majority of Crx+ cells at the apical side incorporated EdU at 48 hpf, indicating that most PRpr are cycling at this developmental stage (Supplementary Figure 1 G). To test whether indeed all PRpr undergo an additional division, we performed photoconversion experiments using a Crx:H2B-Dendra construct at 42 hpf as shown in Figure 1F. 24 hours after photoconversion, 18/19 PRpr (N = 16 embryos) divided (Figure 1 F), while 1/19 did not divide. In 16 out of these 18 PRpr divisions, two PRs were produced (Figure 1 F, G), while in 2/18 cases three or four PRs were produced. However, in these cases we cannot exclude that initially two PRpr were photoconverted. These results indicate that PRs arise from immature committed precursors (PRpr) that divide once to produce two PRs (Figure 1 G).

While this data shows that each Atoh7+ progenitor division gives rise to one PRpr throughout the neurogenic window, the sister cell of the PRpr acquired a different fate. It either became a retinal ganglion cell (RGC)^36, 37^ (Figure 1 C, Supplementary Figures 1 A, B, Video 1) or an inhibitory Neuron (IN), i.e., a horizontal cell precursor^43–45^ (HCpr) (Figure 1D, Supplementary Figure 1 C, D, Video 1) or an amacrine cell ^29, 36^(AC) (Figure 1 E, Supplementary Figure 1 E, F, Video 1), but never a second PRpr or a bipolar cell (BC).

To understand whether and how the distribution of the fates of the PRpr sister cell changed over time, we analysed neuronal fate proportions during two neurogenic windows: between 28 hpf and 42 hpf (from here on referred to as ‘early’) and between 36 and 60 hpf (from here on referred to as ‘late’).

In the early window, 80.8% of divisions (with confidence interval CI = [53.5%, 100%]) produced an RGC, while 19.2% (CI = [0.0%, 46.5%]) produced an inhibitory neuron (ACs or HCs) (Figure 1H). As neurogenesis progressed, the proportion of RGCs decreased (42.9%, CI = [36.0%, 53.7%]) while the proportion of inhibitory neurons increased (57.1%, CI = [46.3%, 64.0%]) (Figure 1H). This indicated that the sister cell fate is acquired with time-dependent probabilities.

While it has been suggested that retinal progenitor competence changes during development, whether this change occurs gradually or more abruptly was still debated^22, 23, 46^. To answer this question, we plotted the time of each progenitor division and its fate outcome for the PRpr sister cell (from here on referred to as ‘*event plots’*). This analysis showed that Atoh7+ progenitors gradually change competence during development (Figure 1 I), switching from a prevalent production of RGCs at early stages of neurogenesis to a progressive decrease of their production and an increased generation of INs. This is consistent with the previously suggested overlapping birth order of retinal neurons^19, 22, 47, 48^.

We conclude that within the same lineage, deterministic and probabilistic fate decisions can co-exist. Atoh7+ progenitor divisions throughout the neurogenic window always generate one PRpr and a sister cell of variable fate. The sister cell can become an RGC or an inhibitory neuron. We never observed a bipolar cell (BC) or a second PRpr. Furthermore, we find that the probabilities of producing RGCs or INs gradually changed over development.

### Bipolar cells arise from the Atoh7 negative sister cell of the Atoh7+ progenitor

We showed that Atoh7+ progenitor divisions can give rise to PRpr, RGC, HC and AC, but we never observed a division producing a BC (Figure 1). This is in line with the previous notion that in zebrafish BCs arise from a different lineage that does not express Atoh7^32^. Analysis of a double transgenic line expressing Atoh7 and Vsx1 (a BC marker^32, 43^) confirmed that BCs do not express Atoh7 (Supplementary Figure 2 A, B). As we previously showed that divisions that produce one Atoh7+ progenitor are asymmetric and also produce one Atoh7-negative progenitor cell (Atoh7-, Supplementary Figure 2 C)^31^, we asked whether BCs arise from this Atoh7-sister progenitor lineage (Supplementary Figure 2 C). Notch inhibition using the gamma-secretase inhibitor LY411575 from 24 hpf is known to induce symmetric division that produce two Atoh7+ progenitors (Supplementary Figure 2 D)^31^ and thus in this condition less bipolar cells should emerge. Inhibiting Notch before neurogenesis onset at 24 hpf indeed resulted in a dramatic reduction of Vsx1+ BCs and an increase of Atoh7+ neurons in the BC layer (Supplementary Figure 2 E). To exclude that Notch inhibition affects BC fate *per se*, Notch was inhibited from 45 hpf onwards, a stage at which BC precursors already emerged^29, 43, 49^. This treatment did not affect the BC population (Supplementary Figure 2 E, bottom panel), showing that BC fate decisions were not generally impaired upon Notch inhibition. Thus, BCs indeed seem to arise from the Atoh7-negative sister cell.

### Perturbing the deterministic fate decision affects overall tissue development and generates non-canonical cell fates

The fact that one PRpr was always produced during the complete neurogenic window made us ask how the outcome of Atoh7+ division would change upon perturbation of this branch of the lineage. To prevent the development of PRprs and consequently PRs, we evaluated target genes resulting from a transcriptomic analysis of 42 hpf retinae, a stage at which photoreceptors are still in their committed precursor state, before terminal division (Supplementary Figure 1 G). These cells already express genes known to be involved in PR development and maturation as *crx*, *prdm1a* and *otx2b*^50–55^ (Supplementary Figure 3 A, B, reference GSE194158 in NCBI). Differential gene expression analysis revealed that only Prdm1a and Crx were significantly enriched in the PRpr population when comparing PRpr to their sister RGC (Supplementary Figure 3 C). While *crx* is linked to PR differentiation and its knockdown leads to PR degeneration^55, 56^, *prdm1a* is involved in PR fate specification^50, 57^. A mouse knockout of *prdm1a* led to a reduction in the number of PRs^50, 58^, without severe consequences on tissue development. Further, our transcriptomics analysis showed that Prdm1a expression is specific to PRprs (Supplementary Figure 3 D). Taking these findings into account, we decided to use a previously established Prdm1a morpholino knockdown approach to interfere specifically with the emergence of PRprs^59, 60^. We found that Prdm1a morphant retinas appeared smaller than controls at 48 hpf (Figure 2 A) and showed more severe microphthalmia by 72 hpf (Figure 2 B, E, N = 12 embryos). While control embryos at 48 hpf feature a layer of Ath5+Crx+ photoreceptors at apical positions, this layer is missing in most of the Prdm1a morphants (Figure 2 A, A’). At 72 hpf a layer starts to form, but it is mostly occupied by Crx- and Zpr1-negative cells, confirming a significant reduction in PR production (Figure 2 B, B’, N = 10 embryos). In extreme cases we found a total depletion of this cell layer (3/10 embryos, Figure 2D). Further, an overall reduction of retinal thickness, mostly due to shrinkage of the outer nuclear layer (ONL) and of the ganglion cell layer (GCL) was observed (Figure 2 C, D, N = 4 embryos (control), 6 embryos (Prdm1a morphant)) as well as a reduction in retinal diameter compared to controls (Figure 2 E, N = 7 embryos (control), 10 embryos (Prdm1a morphant)).

**Figure 2.**
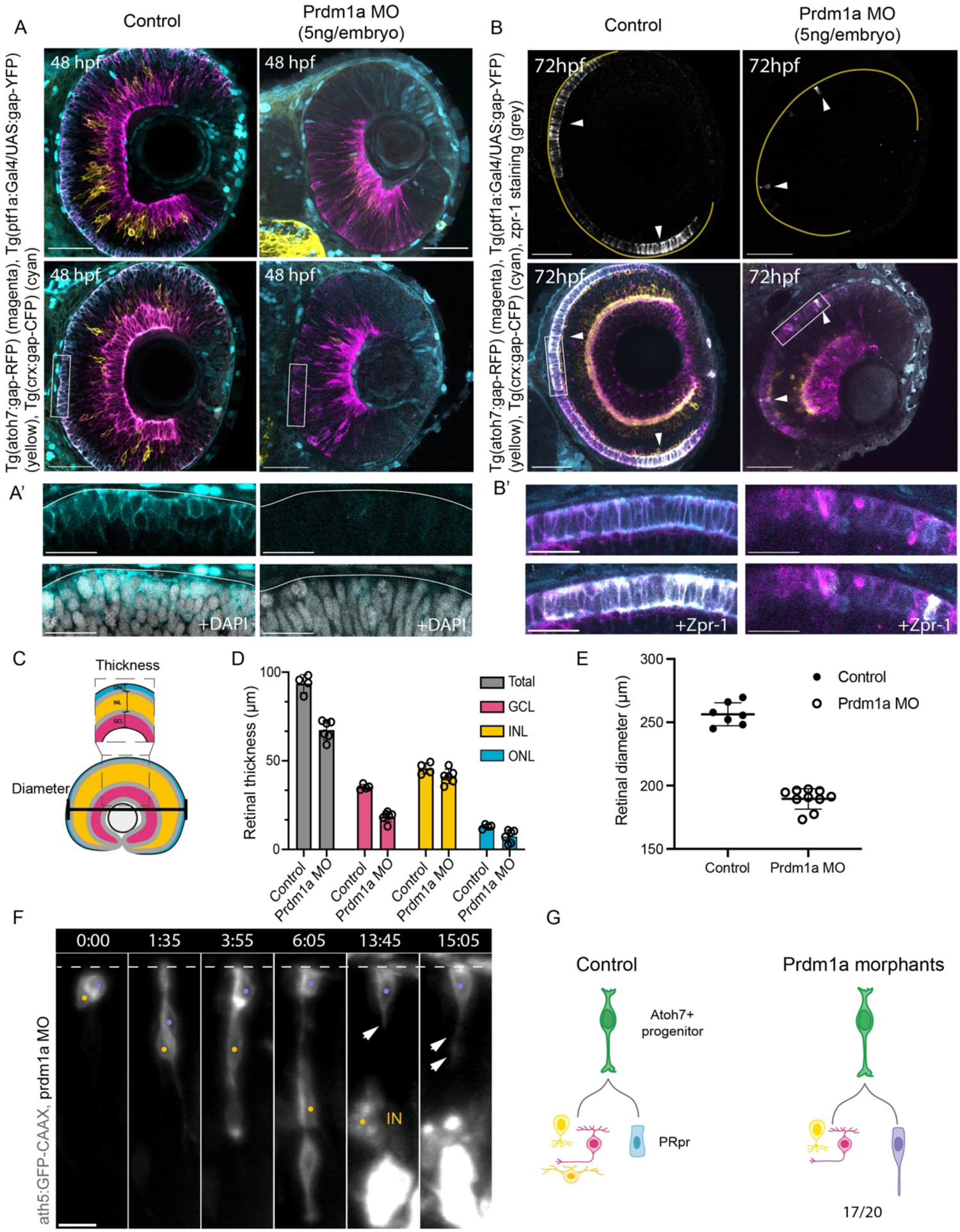
Perturbing the deterministic PRpr fate decision affects lineage and overall tissue development. A) Two examples of retinas at 48 hpf in (left) control and (right) Prdm1a morphant (MO). Atoh7+ cells (magenta), inhibitory neurons (yellow) and photoreceptors (cyan) are labelled. Scale bar 50 µm A’) Close up of Crx (cyan) signal (upper panel) and DAPI (grey, lower panel) from figure A, for controls (left) and Prdm1a morphant (right). Scale bar 20 µm. A) B) Staining for the photoreceptor cell marker zpr-1 at 72 hpf in (left) control and (right) Prdm1a knockdown. Atoh7+ cells (magenta), inhibitory neurons (yellow), photoreceptors (cyan) and zpr-1 (grey). Scale bar 50 µm. B’) Close up of Atoh7 (magenta) and Crx (cyan) signal (upper panel), together with Zpr-1 (grey, lower panel) from figure B, for controls (left) and Prdm1a morphant (right). Scale bar 20 µm. C) Schematics of layer thickness and retinal diameter measurements in the central part of the retina. D) Layer thickness analysis in control and Prdm1a knockdown embryos. N = 4 embryos (control), 6 embryos (Prdm1a morphant). Total thickness comparison: p = 0,0055; GCL comparison, p < 0,0001; INL comparison, ns; ONL comparison, p = 0,0369. Two-way Anova with Bonferroni correction. Mean and SD are indicated, as well as single values. E) Measurements of retinal diameter in control (black dots) and Prdm1a knockdown (empty dots). N = 7 embryos in control, 10 embryos in Prdm1a knockdown, 2 independent experiments. p = 0.0001, Kolmogorov-Smirnov test. Mean and SD are indicated, as well as single values. F) Montage of Atoh7+ progenitor division upon Prdm1a knockdown, generating an IN (yellow dot) and a non-canonical sister cell (violet dot). Dashed line indicates the apical side, arrows indicate the basal process of the sister cell. ath5:GFP-CAAX (Atoh7, grey), scale bar 10 µm. G) Schematic comparison of the outcome of Atoh7+ progenitors in control and Prdm1a morphants.

Together, these results indicated that overall tissue architecture was compromised upon inhibition of PRpr emergence.

We next set out to understand how the outcome of Atoh7+ progenitor divisions changed upon inhibition of PRpr emergence. Possibilities included a fate switch to other neuronal fates, differentiation into a different cell type or progenitors skipping this division completely and directly generating one neuron. 20 divisions Atoh7+ progenitor divisions from three embryos were followed in the early neurogenic window in Prdm1a morphants as established in controls. This revealed that in 2/20 cases Atoh7+ cells did not divide but differentiated directly, one generating an RGC (Supplementary Figure 3 E) and one generating an inhibitory neuron (AC, Supplementary Figure 3 F). One division produced an RGC and a PRpr as seen in controls. Most divisions (17/20), however, generated an RGC or an inhibitory neuron as seen in controls, and a sister cell that initially showed PRpr-like unipolar morphology during basal migration (Figure 2 F, Video 2)^34^. Afterwards, this cell extruded a dynamic basal process and positioned itself apically (Figure 2 F-G) a phenomenon never observed in controls. Upon >15h of imaging, these cells did not acquire any previously observed neuronal morphology and their basal process did not establish a basal attachment (Figure 2 F, compare with Figure 1 B-E).

Thus, interference with the deterministic part of the lineage resulted in lineage and tissue morphology defects, possibly due to the production of a non-canonical sister cell of unknown state from Atoh7+ progenitor divisions. These results also indicate that PRprs are committed to the PR fate and cannot revert to a multipotent state to produce a different, functional cell type when the PR fate cannot be acquired.

### Lineage topology is not affected by interference with the probabilistic lineage branch

Interference with the emergence of PRprs led to severe defects in lineage progression. To understand whether similar constraints in progenitor competence and potency occurred in the probabilistic branch of the lineage, we used knockdown approaches to suppress the emergence of RGCs, inhibitory neurons or both. Particularly, we used established morpholinos against the two bHLH pro-neural transcription factors Atoh7 (prevents RGC fate specification^61^) and Ptf1a (prevents amacrine and horizonal cell fates^33^). Despite the absence of one or two retinal populations, retinal thickness did not significantly change in Atoh7 and Ptf1a morphants (Figure 3 A, B, Table 1). This is consistent with the minimal defects previously observed at the tissue level in Atoh7 and Ptf1a morphants^62, 63^. Analysis of the thickness of different layers showed that the observed conserved retinal size could results from size changes at the level of the different neuronal layers: in Atoh7 morphants, a significant reduction in GCL thickness was counteracted by a thickness increase of the inner nuclear layer (INL) and ONL (mean, SD and p values for the different conditions in Table 1); in Ptf1a morphants, the absence of ACs and HCs lead to a reduction in the thickness of the INL but showed increased GCL and ONL thickness (Figure 3 B, Table 1).

**Figure 3:**
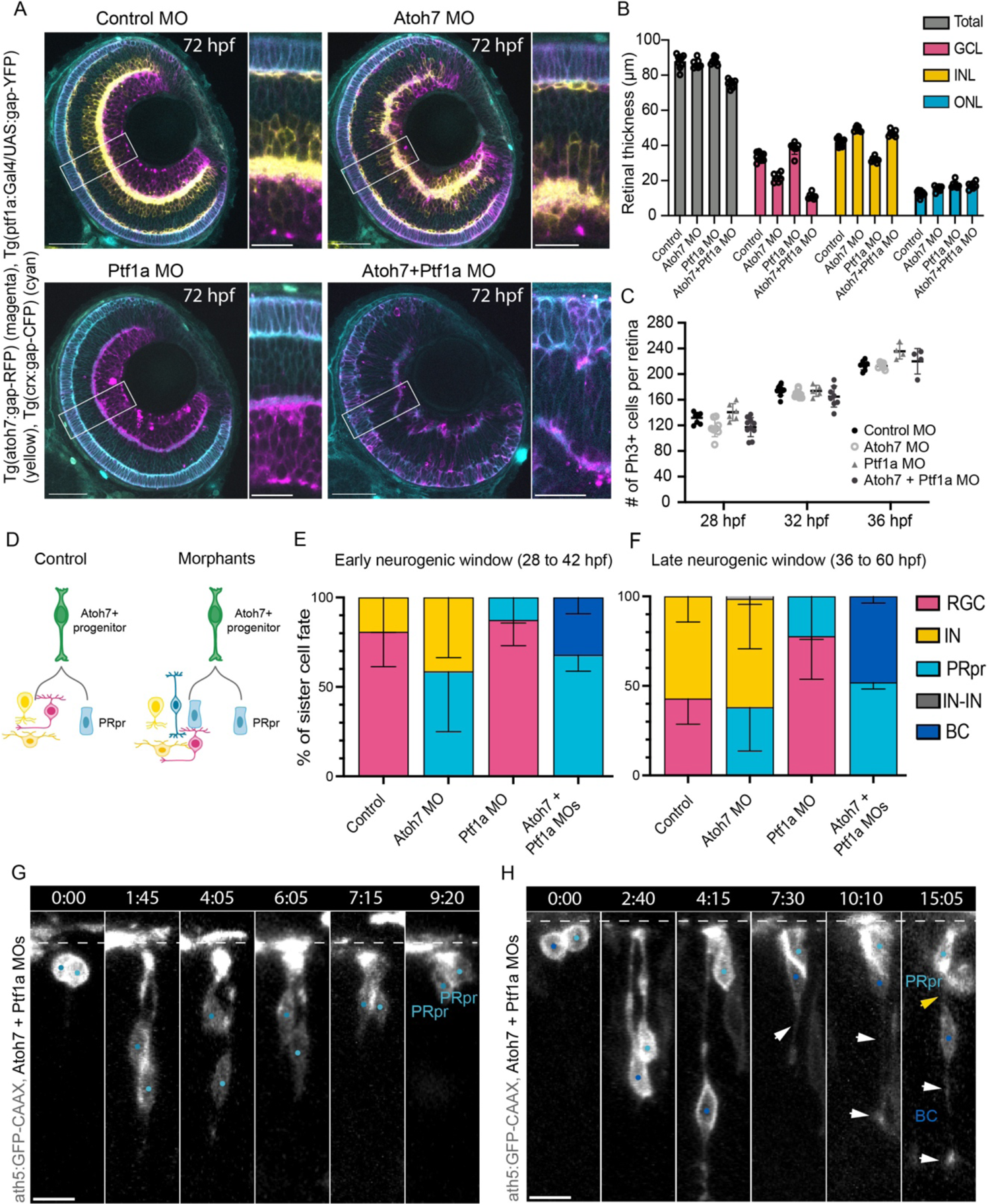
Probabilistic lineage branch shows flexibility upon interference. A) Retina at 72 hpf in (top left) control, (top right) Atoh7 knockdown, (bottom left) Ptf1a knockdown and (bottom right) Atoh7 and Ptf1a knockdown. Atoh7+ cells (magenta), inhibitory neurons (yellow) and photoreceptors (cyan). Scale bar 50 µm, 20 µm in close-up panels. B) Layer thickness analysis in control and morphant embryos. N = 9 embryos (control), 7 embryos (Atoh7 morphant), 7 embryos (Ptf1a morphant), 7 embryos (Atoh7+Ptf1a morphant). p values for thickness measurements are found in Table 1. Mixed effects analysis with Bonferroni correction. Mean and SD are indicated, as well as single values. C) Number of PH3+ cells per retina in control and morphant conditions at 28, 32 and 36 hpf. N = 4 to 10 embryos per condition, mixed effects analysis with Dunnet’s correction. All comparisons are statistically non-significant. Mean and SD are indicated, as well as single values. D) Schematic comparison of the outcome of Atoh7+ progenitor divisions in control and morphants. E) Proportions of PRpr sister cell fates during early neurogenesis. Mean and 95% CI are indicated. F) Proportions of PRpr sister cell fates during late neurogenesis. Mean and 95% CI are indicated. G) Montage of Atoh7+ progenitor division upon Atoh7 and Ptf1a knockdown, generating two PRpr (cyan dots). Dashed line indicates the apical side. ath5:GFP-CAAX (Atoh7, grey), scale bar 10 µm. H) Montage of Atoh7+ progenitor division upon Atoh7 and Ptf1a knockdown, generating a BC (blue dot) and a PRpr (cyan dot). Dashed line labels the apical side, yellow arrow points at BC apical process, white arrows point at BC basal process. ath5:GFP-CAAX (Atoh7, grey), scale bar 10 µm.

**Table 1.**
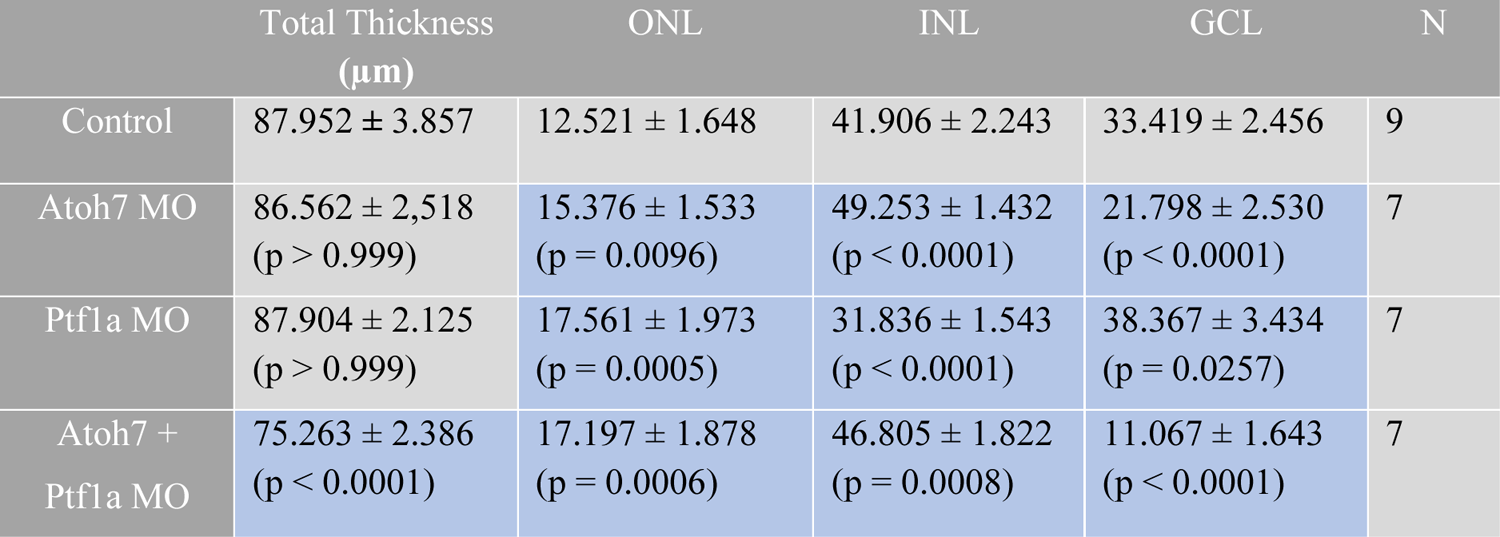
Layer thickness measurements for control, Atoh7-MO, Ptf1a-MO and Atoh7+Ptf1a-MO conditions. Mean and SD are shown for each measurement. p values of comparison with control are shown for each measurement for each morphant condition. N refers to the number of embryos analysed. Blue boxes indicate measurement that are statistically significantly different from the control condition, grey boxes indicate not statistically significant comparisons. Mixed effect analysis with Bonferroni correction for multiple comparisons.

Even in double Atoh7/Ptf1a morphants, in which RGCs, ACs and HCs were missing, only a minor reduction in retinal thickness was observed, mainly due to shrinkage of the GCL (Figure 3 B, Table 1), as previously proposed^62^. This showed that upon interference with the probabilistic branch of the linage, retinal thickness and lamination were generally maintained even in the absence of one or more neuronal cell types. This apparent tissue size robustness is accompanied by changes in the thickness of single layers, suggesting changes in the proportions of neuronal output of progenitor divisions.

One previously suggested possibility that could explain the changes in neuronal proportions was that, in the absence of pro-neural factors as Atoh7, progenitors would go through an additional round of cell division to then produce late-born neurons^23, 24, 64^. If this was the case, a higher number of progenitors dividing at the onset of neurogenesis would be expected. This would lead to changes in the number of progenitor divisions and in the lineage branching structure, here referred to as *lineage topology*. To understand whether lineage topology changed upon knockdown of the pro-neural transcription factors Atoh7 and Ptf1a, we used the mitotic marker PH3 and assessed the number of PH3+ cells at 28, 32 and 36 hpf in single Atoh7 and Ptf1a morphants and in Atoh7/Ptf1a double morphants. Interestingly, no major difference in the number of mitotic progenitors was observed in the four conditions over development (Figure 3 C). We then followed single Atoh7+ progenitors divisions in the three morphant conditions (the Atoh7 morpholino blocks only the expression of the protein, not the reporter) (n = 290 divisions, N = 24 embryos) and found that these divisions never produced an additional Atoh7+ progenitor (Figure 3 D), indicating that the absence of the pro-neural factors Atoh7 and Ptf1a did not delay neuronal production. Further, plotting the developmental time of each neurogenic division showed that the temporal distribution of neurogenic divisions was similar in control and single morphant conditions, with only a slight delay in the double morphants (Supplementary Figure 4 A). This suggested that knockdown of pro-neural genes neither affects the topology of progenitor lineages nor the timing of neurogenic entry.

### Atoh7+ progenitors restrict competence without losing potency

To understand how the outcome of Atoh7+ divisions changed upon interference with the likelihoods of different neuronal fates, we analysed the possibilities and constraints of Atoh7+ progenitors fate decisions in different morphants. Interestingly, independently of the morphant condition (n = 290 divisions, N = 24 embryos), all Atoh7+ progenitor divisions nevertheless produced one PRpr and a neuronal sister cell (Figure 3 D), as seen in controls. Thus, while the deterministic PRpr fate remained unchanged, the sister cell acquired one of the other available fates with different probabilities (Figures 3D-F). This is consistent with the previously proposed ‘fate switch’ hypothesis^33, 63^. Differently from what was seen in controls however, symmetric divisions producing two PRpr (and consequently four PRs) occurred in all morphant conditions (Figure 3 D, G, Video 3). This could explain the increased thickness of the PR layer observed in all morphants (Figure 3 B). Furthermore, in double morphants in which neither RGC nor IN fates were available, divisions producing one PRpr and one BC (Figure 3 D, H, Video 3) appeared. As BCs usually do not emerge from Atoh7+ progenitor divisions (Figure 1 and Supplementary Figure 2), these results indicated that Atoh7+ progenitors generally have the potency to produce BCs but are not competent to generate them when Atoh7 and Ptf1a are expressed. The broad spectrum of possibilities for Atoh7+ progenitors fate decisions further suggested that during normal development Atoh7+ progenitors remain multipotent, while their competence is constrained.

In controls, the likelihood that the probabilistic branch of Atoh7+ progenitor divisions acquired certain neuronal fates changed over development. RGCs were more likely produced at early stages while inhibitory neurons became more prevalent later (Figure 1 H, I). To understand whether and to what extent these proportions were affected in the different morphant conditions, the outcome of Atoh7+ progenitor divisions was analysed over time. In Atoh7 morphants (n = 98 divisions, N = 8 embryos), we observed an increase in the proportions of division giving rise to IN, from 19.2% in control to 41.4% (CI = [15.9%, 56.8%]), Figure 3E) at early developmental stages. In addition, and unobserved in controls, divisions giving rise to a second PRpr (58.586%, CI = [43.2%, 84.1%], Figure 3 E, G) appeared. During later neurogenesis stages, the proportions of divisions producing an IN remained invariant compared to controls (57.1% in control vs 60.6%, with CI = [42.0%, 78.8%] in Atoh7 morphant, Figure 3F) but the proportions of divisions that would have produced RGCs in controls, now produced a second PRpr (38.1%, with CI = [21.2%, 53.2%]) (Figure 3F).

In Ptf1a morphants, (n = 80 divisions, N = 9 embryos) the proportion of divisions that would have produced an IN in controls produced a second PRpr (12.6% with CI = [1.9%, 19.7%]) at early stages (Figure 3 E), while the production of RGCs remained invariant compared to controls (80.8% in control vs 87.4% with CI = [80.3%, 98.1%] in Ptf1a morphant, Figure 3 E). However, at later stages the proportions of RGCs increased with respect to controls (from 42.9% to 77.7% with CI = [67.2%, 96.1%]), indicating that the window of RGC formation is extended in this condition (Figure 3 F). This prolonged production of RGCs could also explain the increased RGC layer thickness observed (Figure 3 B). In double morphants (n = 112 divisions, N = 7 embryos), bipolar cells were produced throughout the entire neurogenic window (Figure 3 F, H). This is interesting for two reasons, a) they were never produced by the Atoh7 lineage in control embryos (Figure 1, Supplementary Figure 2), and b) they usually start emerging only at later developmental stages from 45 hpf^43^.

Together, these results indicated that Atoh7+ progenitor competence and potency are extended in morphant conditions when compared to controls. Our data showed that these progenitors can in principle generate all retinal neuronal fates and that these fates can be produced even outside their canonical temporal window. These experiments also reinforced the finding that deterministic and probabilistic fate decisions co-exist in the same lineage, as the deterministic PRpr production was not altered in any of these morphant conditions.

### A simple theoretical Gene Regulatory Network can explain temporal changes in progenitor competence

Our transcription factor (TF) knockdown experiments revealed an unexpectedly broad spectrum of possibilities for fate decisions in the probabilistic branch of the lineage. Previous studies suggested that interactions between the pro-neural TFs in the retina in the form of gene regulatory networks (GRN) influence the proportions of different neuronal fates within a clone^32, 33^. We thus set out to use the lineage data obtained from the knockdown experiments to develop a minimalistic stochastic model capturing possible interactions between Atoh7, Ptf1a, and Prdm1a. This model used the morphant data to probe whether a simple GRN based on interactions between these transcription factors could explain the temporal changes in progenitors’ competence observed in control conditions.

Event plots were generated for Atoh7, Ptf1a and Atoh7/Ptf1a morphants conditions (Figure 4 A, Supplementary Figure 4B) and the distribution of fate outcomes was renormalised time-point wise by the distribution of all neurogenic division events. This allowed us to calculate the probability of different neuronal fates for a given division occurring at a certain time, which revealed the fate share changes over development (Figure 4 B). We found that when Ptf1a is knocked down, the likelihood of RGC production remained constant over time. Conversely, the production of INs was only slightly changed upon Atoh7 knockdown. We thus hypothesized that, as previously proposed by Jusuf et al.^33^, Ptf1a could have an inhibitory effect on Atoh7, while Atoh7 had no inhibitory effect on Ptf1a. As bipolar cells emerged from Atoh7+ divisions only when both Atoh7 and Ptf1a were depleted (Figure 3 E, F, H), we further hypothesized that expression of either of these genes was sufficient to inhibit bipolar cell transcription factors such as Vsx1. The fact that a second PRpr is generated in all morphant scenarios (Figure 3 E-G) made us add the assumption that PR genes were inhibited by the expression of Atoh7 and/or Ptf1a.

**Figure 4:**
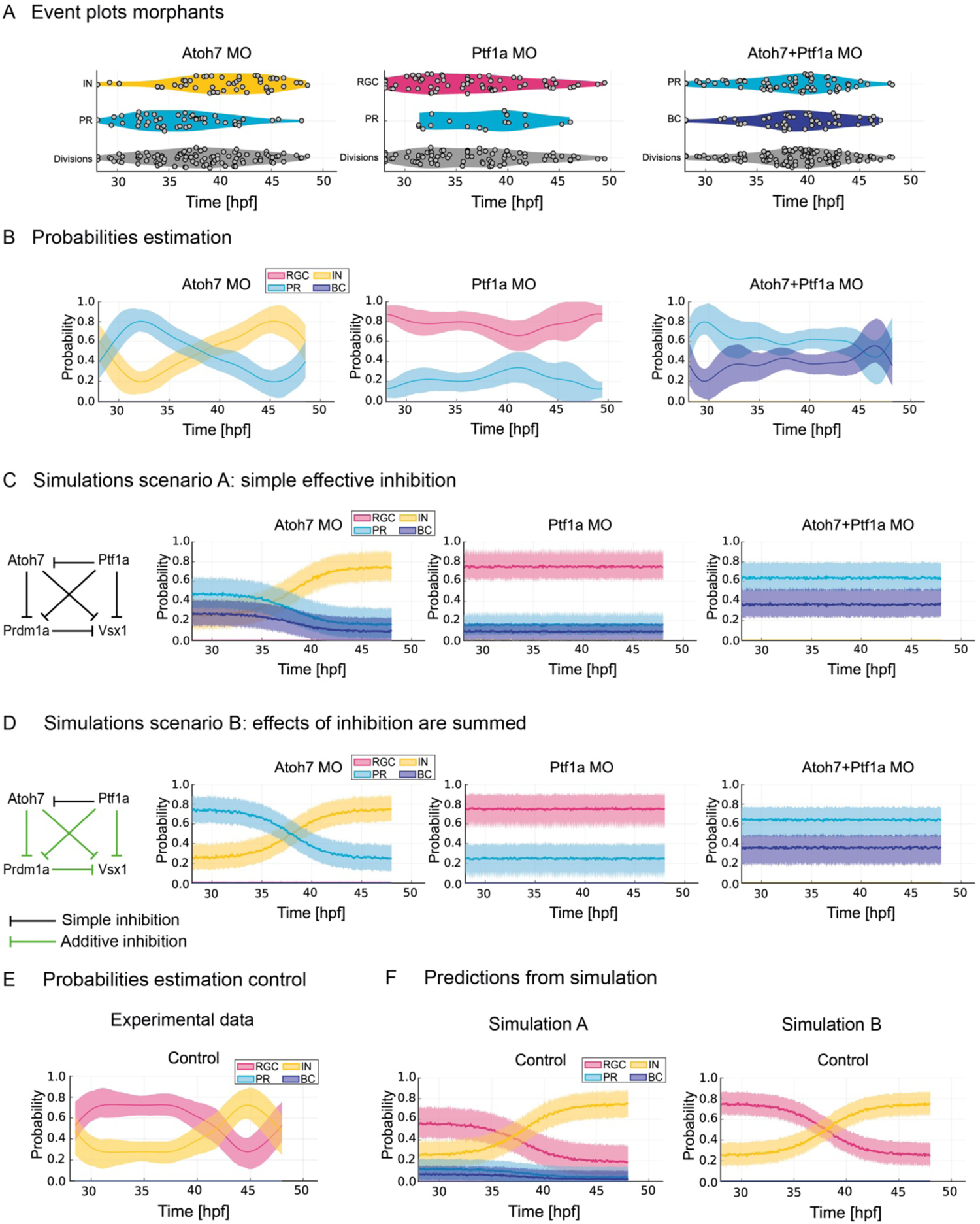
A simple theoretical Gene Regulatory Network reproduces temporal changes in progenitor competence. A) Distribution of each Atoh7+ division in time (Event plots) from 28 hpf in Atoh7 morphants, Ptf1a morphants and double Atoh7/Ptf1a morphants. Coloured violin plots indicate different fates, grey violin plots show all neurogenic divisions analysed. Time in hpf. B) Probability estimation from lineage data in (A) for each fate outcome at the time at which an Atoh7+ division occurs in Atoh7 morphants, Ptf1a morphants and double Atoh7/Ptf1a morphants. Time in hpf. Mean (dark line) and 95% confidence interval (thick transparent stripe) are plotted. C) Simulation scenario A run using a simple GRN tested based on the hypothesis formulated based on the probability estimation plots in (B). Mean (dark line) and 95% confidence interval (thick transparent stripe) are plotted. D) Simulation scenario B run using a simple GRN in which inhibitory effects of two or more TFs are summed (green lines), tested based on the hypothesis formulated based on the probability estimation plots in (B). Mean (dark line) and 95% confidence interval (thick transparent stripe) are plotted. E) Probability estimation from lineage data in Figure 1I for each fate outcome at the time at which an Atoh7+ division occurs in controls. Time in hpf. Mean (dark line) and 95% confidence interval (thick transparent stripe) are plotted. F) Prediction of fate shares in the control situation using the Simulation scenario A (left) and the Simulation scenario B (right). Mean (dark line) and 95% confidence interval (thick transparent stripe) are plotted.

As the knockdown of either Atoh7 or Ptf1a or both did not alter overall lineage topology (Figure 3 C, D, Supplementary Figure 4 A), but only affected fate decisions at the terminal point of the lineage, our model neglected the number of progenitor divisions that occurred before neurogenesis onset.

All above stated assumptions were implemented in a phenomenological stochastic model to test whether a hypothetical GRN based on the morphant lineage data could predict the temporal dynamics of fate decisions observed in control conditions. In the model, noise came from the base levels of transcription factor expression^65–68^ and the inhibitions were modelled considering the presence of a threshold level of expression of each factor. If the expression level of a TF was above the threshold, then the inhibition took place on its target TFs. To model the time-evolution of fate probabilities of experimental data, we considered that TF levels change during development, as previously shown^69^. Interestingly, assuming that only Ptf1a increased it expression over time^69^ was sufficient to completely reproduce the temporal changes of fate probabilities (see Methods for details).

By considering this GRN and implementing a simple effective inhibition (Figure 4 C), the simulation recapitulated the double morphant condition well (compare Figure 4 C right panel with 4 B right panel) but failed to recapitulate the fate outcome and the changes in fate shares over time in the single morphant conditions (compare Figure 4 C middle and left panels with 4 B middle and left panels). In particular, the simulation showed the presence of divisions generating a BC also in single morphants for Atoh7 and Ptf1a, an outcome not observed in our experiments. When generating a prediction for the control condition using this model GRN, the simulation predicted that BCs and a second PRpr could be generated (Figure 4 F, Simulation A). However, this was not found in our experimental dataset (Figure 4 E, Figure 1, Supplementary Figure 2).

When the inhibition mode of the model was amended to a *joint inhibition*, i.e., considering all inhibitors of a given target to sum their levels to reach the threshold, while keeping all other parameters constant (Figure 4 D), the model faithfully recapitulated the fate shares and the temporal dynamics of fate decisions in all morphant conditions (compare probabilities estimation in Figure 4 D with Figure 4 B). Further, and most importantly, the simulation based on this second GRN also correctly predicted the temporal changes of fate shares in the control condition (Figure 4 F, Simulation B), including the lack of bipolar cells production and the absence of symmetric divisions generating two PRpr (compare Figure 4 E with Figure 4 F Simulation B).

Thus, a simple GRN based on the fate probabilities observed in our knockdown experiments can recapitulate the temporal changes in neuronal proportions in control retinas. This suggests that seemingly complex temporal shifts in progenitor competence observed during neurogenesis could originate from a simple gene regulatory pattern in a noisy environment. In principle, this means that such a simple GRN could be sufficient to achieve the observed fate shares during development from multipotent Atoh7+ progenitors.

## Discussion

In this study we investigated the possibilities and constraints of Atoh7+ progenitor neurogenic fate decisions during zebrafish retinal development.

Analysis of the output of single progenitor divisions revealed that two different types of fate decisions, deterministic and probabilistic, co-exist in the same lineage and that these are characterized by different probability distributions. We further find that upon interference with these different fate decisions, the lineage responds differently depending on which branch is affected. Upon interference with PRpr emergence, the deterministic lineage branch does not produce physiologically observed fates. This contrasts with what is observed upon interference with the more probabilistic lineage branch, where a broader spectrum of fate outcomes was observed even compared to the control scenario.

Stochastic modelling of the fate outcome of the probabilistic branch in perturbed conditions let us propose a GRN that predicts the temporal changes in progenitor competence in controls. This simple GRN was based on the observed changes in fate probabilities and explained the temporal production of different retinal neurons in the right proportions during development. It is thus tempting to speculate that the “fate plasticity” unravelled in this study could be important for robust tissue development.

To date, fate decisions in the vertebrate CNS have mostly been investigated at the clonal or the population level. Depending on the techniques used and interpretation of the data, fate decisions were considered either mostly driven by stochasticity^23–26^ or by deterministically encoded programs in each progenitor cell ^22, 70–72^. Our quantitative analyses of single progenitor divisions and their neuronal output over development shows that both deterministic and stochastic fate decision co-exist in the same progenitor lineage. In the case of the Atoh7+ progenitors, PRpr are always and continuously produced during development in one branch of the lineage, while the sister cell acquires different fates with time-dependent probabilities. Our analysis revealed stereotypic lineage patterns among the general stochasticity in fate decisions: while neuronal fate decisions in one branch of the lineage follow a probability distribution that changes over time, the other branch always and reliably produces one PRpr and consequently two PRs. This adds to the previously proposed stochastic model, in which all neuronal fates are decided according to a set of different probabilities^23, 24^.

We further found that in controls the Atoh7+ progenitors always produced neurons and committed precursors, but never a self-renewing Atoh7+ progenitor (Figure 1). This is different from the Drosophila CNS and the vertebrate neocortex, where asymmetric divisions produce one neurogenic cell and one renewing progenitor or Radial Glia^73, 74^. These renewing progenitors change competence during development, producing different neuronal types in each round of division. We here show that in the zebrafish retina neuronal fate decisions occur at the terminal tips of the lineage. Furthermore, our interference experiments disclosed that in principle Atoh7+ progenitors have the potency to produce all neuronal types (Figure 1, Figure 3) and that their competence windows can extend to earlier or later developmental times (Figure 3 E, F). This suggests that different mechanisms regulating the temporal production of different neuronal types could be at play in the zebrafish retina from what is known in Drosophila^9^. It would be interesting to investigate whether the absence of self-renewing divisions and the presence of lineage patterns that consistently generate one committed PR precursor is specific to the zebrafish retina, possibly due to fast embryo development, or whether it is also found in other areas of the CNS including in other species. Fate-restricted progenitors have been found in the mouse retina^75^ and in the brain neocortex^76, 77^ and it would therefore be interesting to investigate whether these progenitors arise from lineage patterns similar to those found in this study. Studies like this would enhance our understanding of similarities between different CNS areas and species and unveil core principles or differences of those of lineage progression in neural systems.

When we probed the possibilities and constraints of the deterministic and probabilistic fate decisions, interference with the deterministic PR fate affected overall tissue integrity and lineage topology. Interestingly, upon interference with the probabilistic fate decision, Atoh7+ progenitors often generated a <u>second</u> PRpr, a fate not observed in controls for the sister cell. This suggests that generating at least one PRpr is also important for retinal integrity. Consequently, the PR fate could be a “default state” for neurogenesis outcome. This is an attractive idea also in evolutionary terms as PR cells are widespread throughout evolution^78^ and are thought to be the first retinal cell types that evolved^78, 79^. It is therefore possible that the other cell types evolved “on top” of a general PR program as proposed by Constance Cepko in 2014 and Detlev Arendt in 2003^22, 78^. Currently, this is however purely speculative.

Interestingly, in contrast to the deterministic branch, interference with the probabilistic branch of the lineage showed a widened spectrum of fate possibilities for Atoh7+ progenitors. Here, neurons that in controls were mainly produced at early stages were also born later. This finding is in line with previous reports in the RP2/sib neuroblast lineage in Drosophila, where it was shown that progenitors exhibit temporal plasticity and can give rise to early lineages at later stages^80^. Furthermore, in double Atoh7/Ptf1a morphants, BCs were seen to arise from the Atoh7 lineage, an outcome never observed in controls. In this case, these usually late-born neurons appeared earlier than normally observed (Figure 3E). This indicates that Atoh7+ progenitors’ can generally produce any neuronal fate at any time, challenging the idea that retinal progenitor cells can give rise to certain fates only during fixed competence windows^22^.

Overall, the fact that the deterministic and probabilistic lineage branches show different fate outcome possibilities suggests that different molecular mechanisms might act to specify the PRpr vs the sister cell fate. Modelling the fate probabilities observed upon TF knockdown revealed a network of TF interactions that follows these rules of fate decisions. This simple GRN (Figure 4D) featuring Atoh7, Ptf1a and Prdm1a was able to recapitulate temporal changes of the probabilities of generating different fates in the morphant conditions and to accurately predict the changes in progenitor competence in control conditions. We are aware that the scenarios that allowed us to induce fate probability changes were rather artificial, as we suppressed one or more neuronal populations. However, the fact that the lineage reacted to such perturbations with an unexpected broad spectrum of fate possibilities makes us speculate that such lineage flexibility could underlie the presence of plasticity in fate decisions during normal development. As modelling the changes in fate probabilities predicted the temporal changes in progenitors fate decisions in the control scenario, it is possible that fate plasticity could exist to ensure progenitors competence changes during development. However, it is important to note that the model presented here is only one possible scenario. Further, this model can currently not clarify whether the included TFs act directly or indirectly, nor whether further TFs are involved. Thus, additional investigations involving ChiP-sequencing and timely-controlled overexpression studies are needed.

In contrast to the probabilistic fate decisions, the deterministic PR fate decision does not seem to rely on the same GRN, as interference with Prdm1a expression does not result in a fate switch (Figure 2 F, G). This suggests that the PR fate might be acquired through a different mechanism. As a PRpr is reproducibly produced from each Atoh7+ division, one possibility is that fate determinants specifying the PR fate are asymmetrically inherited by the cell that becomes a PRpr. This is a widespread strategy employed by different systems in which asymmetric fate decisions occur, like in the Drosophila neuroblast^9, 73^ or in the mouse retina^81–83^. Transcriptomics analysis combined with advanced live imaging techniques will be needed to find key regulators of this fate decision.

Overall, our study contributes to the understanding of possibilities in lineage decisions in the retina and revealed different degrees of flexibility for different types of fate decisions. Such studies combined with theoretical modelling are important to appreciate and interpret the clonal and molecular data available and thereby understand the core principles of reproducible organ formation.

## Materials and methods

### Zebrafish husbandry

Wild-type zebrafish were bred and maintained at 26°C. Embryos used for experimental work were raised at 21°C, 28.5°C, or 32°C in E3 medium supplemented with 0.2 mM 1-phenyl-2-thiourea (PTU, Sigma-Aldrich) from 8 hours post fertilization (hpf) to prevent pigmentation. Medium and PTU were changed daily. Animals were staged in hpf according to Kimmel et al^84^. All animal work was performed in accordance with European Union directive 2010/63/EU, as well as the German Animal Welfare Act, and in accordance with and the Portuguese legislation (Decreto-Lei n° 113/2013).

### Transgenic lines

Tg(atoh7:gap-GFP) and Tg(atoh7:gap-RFP) zebrafish transgenic lines were used to identify Atoh7+ progenitors and Atoh7+ neurons^37^. The Tg(crx:gap-CFP) line was used to visualize PRs and PRpr^63^. To visualize all different neurons and for transcriptomics, the triple transgenic line Tg(crx:gap-CFP), Tg(atoh7:gap-RFP) and Tg(ptf1a:Gal4/UAS:gap-YFP)^63^ was used.

### DNA injections

DNA was injected at 1 cell stage to mosaically label progenitors and neurons in the retina. 1 nl of ath5:GFP-CAAX^36, 37^ plasmid was injected in each wild type or Tg(atoh7:gap-RFP) embryo in controls and in morphant conditions to label Atoh7+ progenitor cells and their progeny.

### Photoconversion

The Crx:H2B-Dendra construct used to label crx+ cells for photoconversion was assembled using Gateway cloning (Thermo Fisher Scientific) based on the Tol2 kit.

The pME H2B-Dendra was combined with the 5’ entry clone containing the Crx promoter (a kind gift from Rachel Wong) intro the destination vector pTol2+pA+cmlc:eGFP R4-R3^85^. 15-30 ng/µl of the plasmid were co-injected with 2 ng of p53 MO in one-cell stage embryos. Embryos were raised until 42 hpf, mounted in agarose in 35mm glass-bottom petri dishes (Greiner Bio-One) and imaged with spinning-disk confocal. 2-3 cells per embryos were photoconverted. Isolated cells expressing Crx:H2BDendra were chosen for photoconversion according to their proximity to the apical side. After photoconversion, embryos were taken out of agarose and grown for 24 hours in a 12-well plate in E3 medium supplemented with 0.2 mM 1-phenyl-2-thiourea. After 24 hours, embryos were mounted in agarose again for imaging and photoconverted cells were assessed.

### FACS sorting for transcriptomics

For RNAseq of PRprs, RGCs and INs, retinas from a triple transgenic line containing cell-type specific reporters [Tg(crx:gap-CFP), Tg(atoh7:gap-RFP), Tg(ptf1a:Gal4/UAS:gap-YFP)]^63^ were dissociated mechanically. FACS was performed using FacsAria Fusion. The gates were adjusted for autofluorescence/background fluorescence using single transgenic and wild-type embryos. A minimum of 10.000 live cells were sorted per experiment from a pool of 25 retinas. Different neurons were sorted thanks to the expression of a different combination of markers (Supplementary Figure 3A). For transcriptome analysis, 500 cells were sorted per population from a pool of 25 retinas directly into the buffer for extraction.

### RNAseq of FACS sorted zebrafish retinal neurons

For each of the five biological replicates, RNA extraction of FACS sorted PRpr, RGC and INs was performed according to previously published protocol^86^ by the Deep Sequencing Facility at the Genome Center of the Technische Universität Dresden. Data and methodological details (reverse transcription, cDNA amplification, library preparation, sequencing, and data processing) are accessible through GEO series accession number GSE194158 at NCBI.

### Morpholino experiments

To knock down specific genes, the following amounts of morpholinos were injected per embryo into the yolk at 1-cell stage: 2 ng p53 MO, 5’-GCGCCATTGCTTTGCAAGAATTG-3’ (Gene Tools, ^87^); 4 ng Atoh7 MO, 5’-TTCATGGCTCTTCAAAAAAGTCTCC-3’ (Gene Tools,^61^); 10 ng Ptf1a MO1, 5’ - CCAACACAGTGTCCATTTTTTGTGC - 3’ (Gene Tools, ^33^); 10 ng Ptf1a MO2, 5’ - TTGCCCAGTAACAACAATCGCCTAC - 3’ (Gene Tools, ^33^); 5 ng Prdm1a MO, 5’ - TGGTGTCATACCTCTTTGGAGTCTG - 3’ (Gene Tools, ^59, 60^).

### Drug treatments

The Notch inhibitor LY411575 was dissolved in DMSO and used at a concentration of 10 μM. An equal volume of DMSO was used for controls. Embryos were dechorionated and up to 10 embryos were placed in a well of a 24-well plate, then incubated at 28 °C in the dark in E3 medium. The treatment windows are specified in the figure and in the figure legend.

### *In vivo* labelling of proliferative PRprs

To label proliferative PRprs, 48 hpf embryos were incubated at 4°C for 1 h in E3 supplemented with 500 μm of EdU (ClickiT-Alexa 488 fluorophore kit, Invitrogen) in 10% DMSO. After incubation, embryos were washed twice with E3 and immediately fixed overnight in 4% PFA. After antibody staining, incorporated EdU was detected according to manufacturer’s protocol. Embryos were stored in PBS at 4°C until imaging.

### Immunofluorescence

All immunostainings were performed on whole-mount embryos fixed overnight in 4% paraformaldehyde (Sigma-Aldrich) in PBS at 4°C as previously described ^88^.

Embryos were washed five times for 10 min in PBS-T (Triton X-100 in PBS) 0.8%. For permeabilization, embryos were incubated with 0.25% Trypsin-EDTA in PBS on ice for different time depending on the developmental stage (10 min for 24 hpf, 28 hpf and 36 hpf, 12 min for 42 hpf, 15 min for 48 hpf and 17 min for 72 hpf). Embryos were then kept on ice for 30 min in PBS-T 0.8%. Blocking was performed with 10% donkey or goat serum in PBS-T 0.8% for 3 h at room temperature or overnight at 4°C.

Embryos were incubated with the following primary antibodies for 72h at 4°C: GFP (50430-2-AP, Proteintech) 1:100, Histone H3 (phospho S28, ab10543, Abcam) 1:500 and zpr-1 (AB_10013803, zirc) 1:750.

Embryos were then washed 5 times for 30 min with PBS-T 0.8% and then incubated for 48 hours with the appropriate fluorescently labelled secondary antibody (Molecular Probes) at 1:500 and DAPI 1:1000 (Thermo Fisher Scientific). Finally, embryos were washed four times for 15 min with PBS-T 0.8% and stored in PBS at 4°C until imaging.

### Image acquisition

#### Confocal microscopy

Fixed samples were imaged with a laser-scanning microscope (Zeiss LSM 880 Airyscan inverted or Zeiss LSM 980 Airyscan2 inverted, equipped with two PMT and one GaAsP) using the 40×/1.1 C-Apochromat water immersion objective (ZEISS). Samples were mounted in 0.6% agarose in glass-bottom dishes (MatTek Corporation) and imaged at room temperature. Serial sections were acquired every 1 µm with ZEN 2011 (black edition) or Zeiss’s ZEN Blue v3.0.

#### In vivo light sheet fluorescent imaging (LSFM)

Imaging started at 28 hpf for the early neurogenic window experiments and at 36 hpf for the late neurogenic window. Embryos were manually dechorionated and mounted in ∼1mm inner diameter glass capillaries in 0.6% low-melting-point agarose as previously described^38^. The sample chamber was filled with E3 medium containing 0.01% MS-222 (Sigma) and 0.2 mM PTU (Sigma). Imaging was performed on a Zeiss Lightsheet Z.1 microscope equipped with two PCO Edge 4.2 sCMOS cameras (max 30 fps with 2048×2048 pixels - pixel size 6.5 μm) and with a 20x/1,2 Zeiss Plan-Apochromat water-immersion objective. Imaging was performed at 28.5 °C. Z-stacks spanning the entire retinal epithelium (70-100 μm) were acquired with 1 μm optical sectioning every 5 min for 15-24 hours with double-sided illumination mode. The system was operated by the ZEN 3.1 software (black edition).

To confirm the effect of the morpholino injections in live imaging experiments, the absence of the optic nerve was assessed after live imaging of the Atoh7 MO condition. Embryos in which the optic nerve was present were discarded from the analysis. Embryos injected with Ptf1a MOs were grown at 28.5°C until 72 hpf after imaging, fixed in PFA and immunostained against HuC/D (A-21271, Thermo Fisher). Embryos displaying HuC/D staining in the inner nuclear layer were discarded from the analysis.

#### Spinning disk imaging for photoconversion

Photoconversion experiments were performed using an Andor spinning disk confocal microscope composed of Andor IX 83 stand and a CSU-W1 scan head (Yokogawa) with Borealis upgrade, equipped with a DMD Andor Mosaic module and a 405 nm photomanipulation light source. The microscope was operated via the Andor iQ software version 3.6. Embryos were embedded in 0.8% of low melting point agarose in E3 medium supplemented with 314 μg/mL of MS-222 and 0.1 M Hepes (pH 7.25) in 35-mm glass bottom Petri dish (Greiner Bio-One). The dish was filled with E3 supplemented with 120 μg/mL of MS-222. Z-stacks were acquired using Olympus UPLSAPO objective 60x 1.3 SIL and Andor iXon 888 Ultra with Fringe suppression.

### Quantitative analysis

Images from live imaging experiments were cropped and averaged in ZEN Black and/or Fiji^89^ and were corrected for drift using the Fiji plugin “Manual Drift Correction” (https://imagej.net/Manual_drift_correction_plugin) created by Benoit Lombardot (Scientific Computing Facility, Max Planck Institute of Molecular Cell Biology and Genetics, Dresden, Germany).

### Cell fate assignment

To identify the fate of the daughter cells of Atoh7 expressing progenitors, single cells were followed until they reached their final position or until they acquired a distinct morphology (Figure 1B,C). Morphology was assessed by the use of the membrane marker ath5:GFP-CAAX^88^ in combination with the transgenic line Tg(atoh7:gap-RFP). PRpr fate was assigned based on unipolar morphology during migration and columnar morphology after reaching the apical side^34, 42, 43^, their bidirectional migration pattern^34^ and final apical positioning. RGC fate was assigned based on basal position, basal somal translocation^88^ and by the emergence of a basal axon (Figure 1C). Inhibitory neuron (ACs and HCs) fate was assigned based on multipolar migration mode^45, 90^, absence of basal axons and by final positioning (figure 1D-E). Bipolar cell fate was assigned by bipolar morphology and nuclear positioning in the INL^43, 49^ (Figure 3H).

### Analysis of cell migration

To generate the trajectories for each cell type in Figure 1B, cells cropped with ZEN 3.1 (Zeiss) and processed in Fiji as described above. Cells were tracked in 2D in maximum projected sub-stacks by following the centre of the cell body in Fiji using the semi-automated ImageJ plugin MtrackJ ^91^. Tracking started at birth of each cell and ended after the ell reached its final position in the tissue.

### Retinal size measurements

Retinal size measurements were performed manually using the Fiji line tool on 72 hpf retinas from the Tg(ptf1a:Gal4, UAS:YFP; atoh7:gap-RFP; crx:gap-CFP) line, in controls and in all morphant conditions as illustrated in Figure 2C. Retinal diameter was measured on three different z planes per retina, and the average measurement was plotted for each replicate.

Retinal thickness was measured on three different z plane of the central region of the tissue, and the average measurement was plotted for each replicate. The outer nuclear layer (ONL) was assigned as the distance between the apical side and the outer plexiform layer; the inner nuclear layer (INL) as the distance between the inner plexiform layer and the outer plexiform layer; the ganglion cell layer (GCL) was assigned as the layer between the inner plexiform layer and the most basal position of the retina.

### PH3+ cells count

To count the number of PH3+ cells in whole retinas, the Tg(hsp70:H2B-RFP) was used to identify the boundaries of the tissue. Then, stacks covering the whole tissue were acquired using the laser-scanning confocal. Images were imported in Imaris and the mask tool was used to isolate the retinal tissue from the rest of the tissues in the image. The spot detection tool was used to count the number of PH3+ cells with these parameters: diameter = 8 µm (x-y dimensions), PSF modelling = 16 µm. Threshold for detection of the PH3+ nuclei was adjusted manually to ensure that all the PH3+ nuclei were counted but was usually kept around 1800.

### Statistical analysis

All Statistical tests used are indicated in the figure legend, as well as the definitions of error bars. All test used were two-sided and 95% confidence intervals were considered. P values are indicated in the figure legends, as well as sample sizes, or in Table 1 for experiments in Figure 3B. Data were analysed using GraphPad Prism 6 or Python 3. Statistical analysis was performed using GraphPad Prism 6 and Julia 1.7.2. Information about the exact libraries, together with their versions, used for the *Event plots* and for the i*n-silico stochastic model* are detailed in the *Project.toml* and *Manifest.toml* files in the respective Git repositories, as per standard dependency handling procedures of the Julia language:

- https://git.mpi-cbg.de/nerli/lineage-analysis
- https://git.mpi-cbg.de/bianucci/transcription-factors-interactions

### Event plots

*Event plots* were produced from the raw experimental data using a custom data analysis and plotting pipeline developed by the authors, which is available at https://git.mpi-cbg.de/nerli/lineage-analysis. The code comes together with the data files, allowing for an easy and complete replication of the analysis and plots that appear in this work.

The data analysis consists of the following steps:

1. Consider the time at which each division occurs as the time of the fate decision event and categorize these events by the fate acquired by the sister cell of the PRpr.
2. Compute the Kernel Density Estimation (KDE) of the distribution of events over time for each fate type and for the total distribution of divisions.
3. Rescale the KDE of each fate by the KDE of all divisions to obtain an estimation of the probability of acquiring the different fates dependent on the time the division occurs.
4. Use the *bootstrap* resampling method^92^ to obtain uncertainties of the KDEs and of the fate probabilities.

### In-silico stochastic model of cell fate decision

To evaluate the potential of candidate GRNs to reproduce our experimental data of the probabilistic branch of the lineage, we designed and implemented a simple *in-silico* phenomenological model of stochastic fate decisions. The code is available at https://git.mpi-cbg.de/bianucci/transcription-factors-interactions.

Since we know from the lineage analysis data that each division produces one PRpr and one sister cell whose fate is stochastically determined, we modelled only one fate decision per division. Also, as the TFs Atoh7, Ptf1a and Prdm1a are responsible for the specification of the different neuronal fates, we modelled their expression as random variables, with time-dependent expected values. Their expression levels were normalized with respect to an *effective threshold*, i.e., a level at which they start to carry out their inhibitory function and produce a downstream effect.

The simulation of this model relies on drawing N=100 fate decisions for each time point to estimate the shares of different fates, this was then repeated for R=100 times to compute the mean and confidence interval of these shares. The TF levels were modelled as normally distributed, with a time-dependent mean value and a constant standard deviation. Being the threshold set at the arbitrary value of 1.0, in the model we set the mean expression levels of Atoh7, Prdm1a at the values of 1.1 and 1.05 respectively. The mean levels for these TFs were kept constant over time. The mean expression level for Ptf1a was instead time-dependent^69^ and increased according to a logistic function with lower and upper asymptotes at 0.9 and 1.1 respectively.

The standard deviation was constant over time for all TFs and was computed by multiplying a fixed coefficient of variation (CV=0.14) by the time-averaged expression level of each TF. Finally, the GRN was modelled as a decision rule occurring at the terminal branching point of the lineage (Figure 1G), taking the TF levels as input, and producing a fate choice as output. Any inhibition in the GRN was translated into a *precedence* rule, i.e., a TF that inhibits a second one is able, if its level is above the threshold, to determine the acquisition of a certain fate, while the inhibited TF is ignored. The precedence rule for Scenario A (Figure 4C,E) is illustrated by the following diagram:

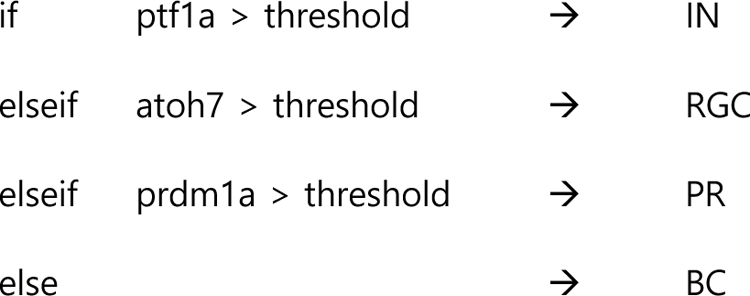

To achieve inhibition of the target TF in additive inhibitions, it was enough that the sum of the levels of the inhibitors involved was above the threshold. The precedence rule in the additive inhibition case (Scenario B, Figure 4D,F) is illustrated by the following diagram:

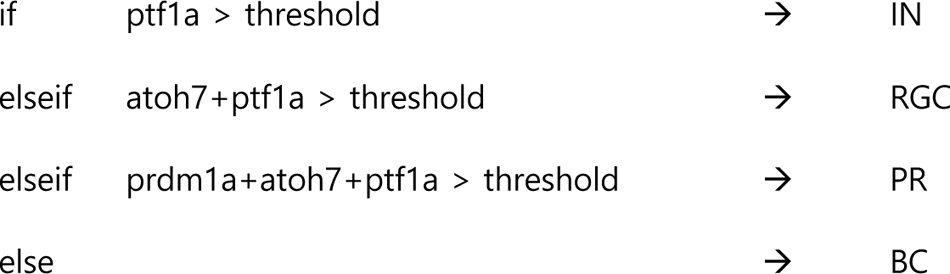

In summary, the fate decision model consists of (1) drawing independent and normally distributed stochastic TF expression levels, (2) applying the decision rule and recording the resulting fate, (3) repeating this stochastic decision for 100 divisions taking place at any given time point and (4) again repeating the whole process for 100 times, creating a synthetic dataset that can then be analysed in the same way as the experimental data.

---

**Supplementary Figure 1.**
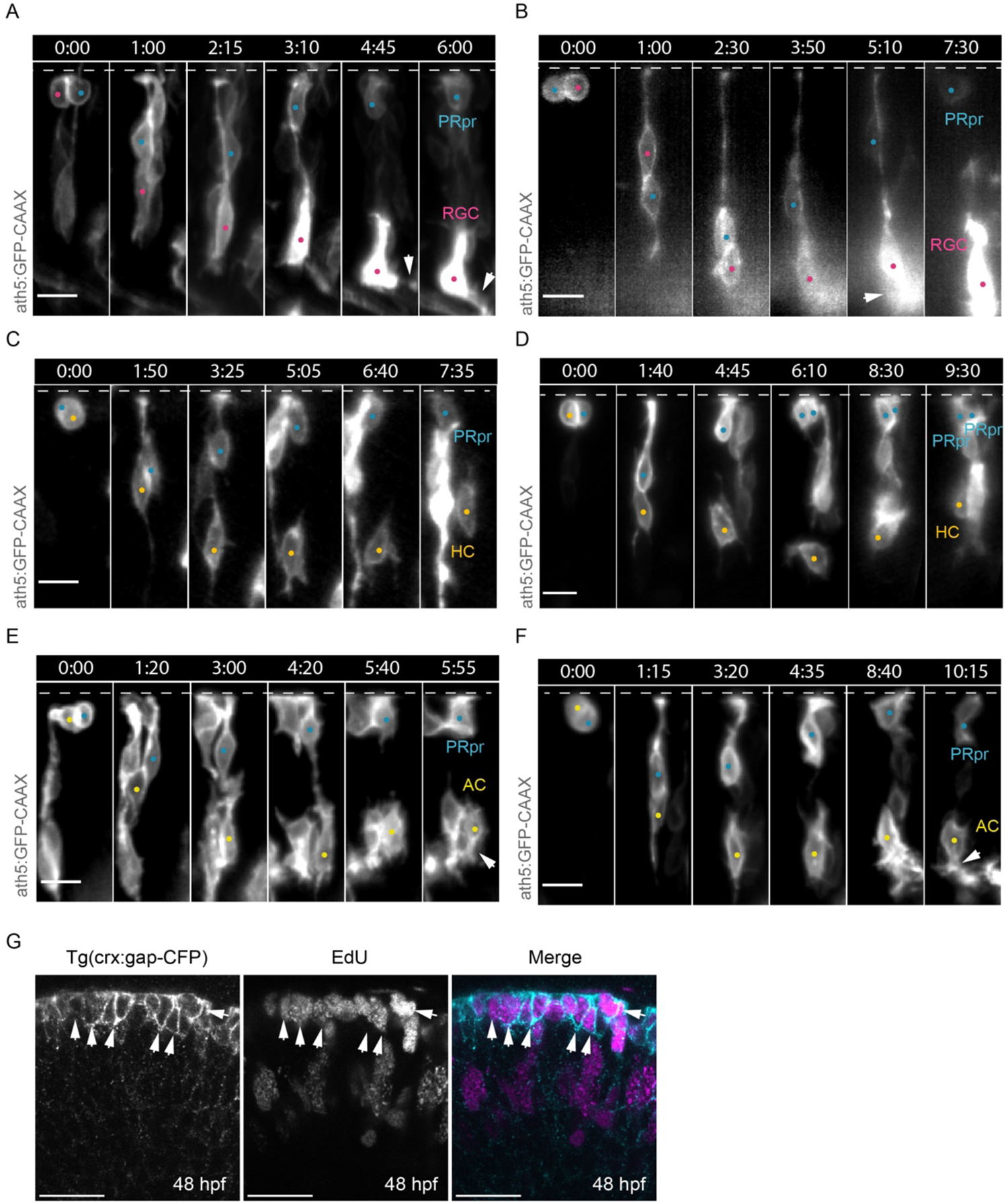
Atoh7+ progenitors divisions producing a PRpr and a sister cell. A) Montage of Atoh7+ progenitor division generating an RGC (magenta dot) and a PRpr (cyan dot). Dashed line indicates the apical side, arrowhead points to RGC axon. ath5:GFP-CAAX (Atoh7, grey), scale bar 10 µm. B) Montage of Atoh7+ progenitor division generating an RGC (magenta dot) and a PRpr (cyan dot). Dashed line indicates the apical side, arrowhead points to RGC axon. ath5:GFP-CAAX (Atoh7, grey), scale bar 10 µm. C) Montage of Atoh7+ progenitor division generating an AC (yellow dot) and a PRpr (cyan dot). Dashed line indicates the apical side, arrowhead points to basal dendrites. ath5:GFP-CAAX (Atoh7, grey), scale bar 10 µm. D) Montage of Atoh7+ progenitor division generating an AC (yellow dot) and a PRpr (cyan dot). Dashed line indicates the apical side, arrowhead points to basal dendrites. ath5:GFP-CAAX (Atoh7, grey), scale bar 10 µm. E) Montage of Atoh7+ progenitor division generating an HC (orange dot) and a PRpr (cyan dot). Dashed line indicates the apical side. ath5:GFP-CAAX (Atoh7, grey), scale bar 10 µm. F) Montage of Atoh7+ progenitor division generating an HC (orange dot) and a PRpr (cyan dot). Dashed line indicates the apical side. ath5:GFP-CAAX (Atoh7, grey), scale bar 10 µm. G) Proliferative status of photoreceptor precursors, labelled by Tg(crx:gap-CFP) (right) and EdU (centre), at 48 hpf. In merged image, Tg(crx:gap-CFP) (cyan) and EdU (magenta). Scale bar, 20 µm, arrowheads point to Crx+/EdU+ PRpr.

**Supplementary Figure 2.**
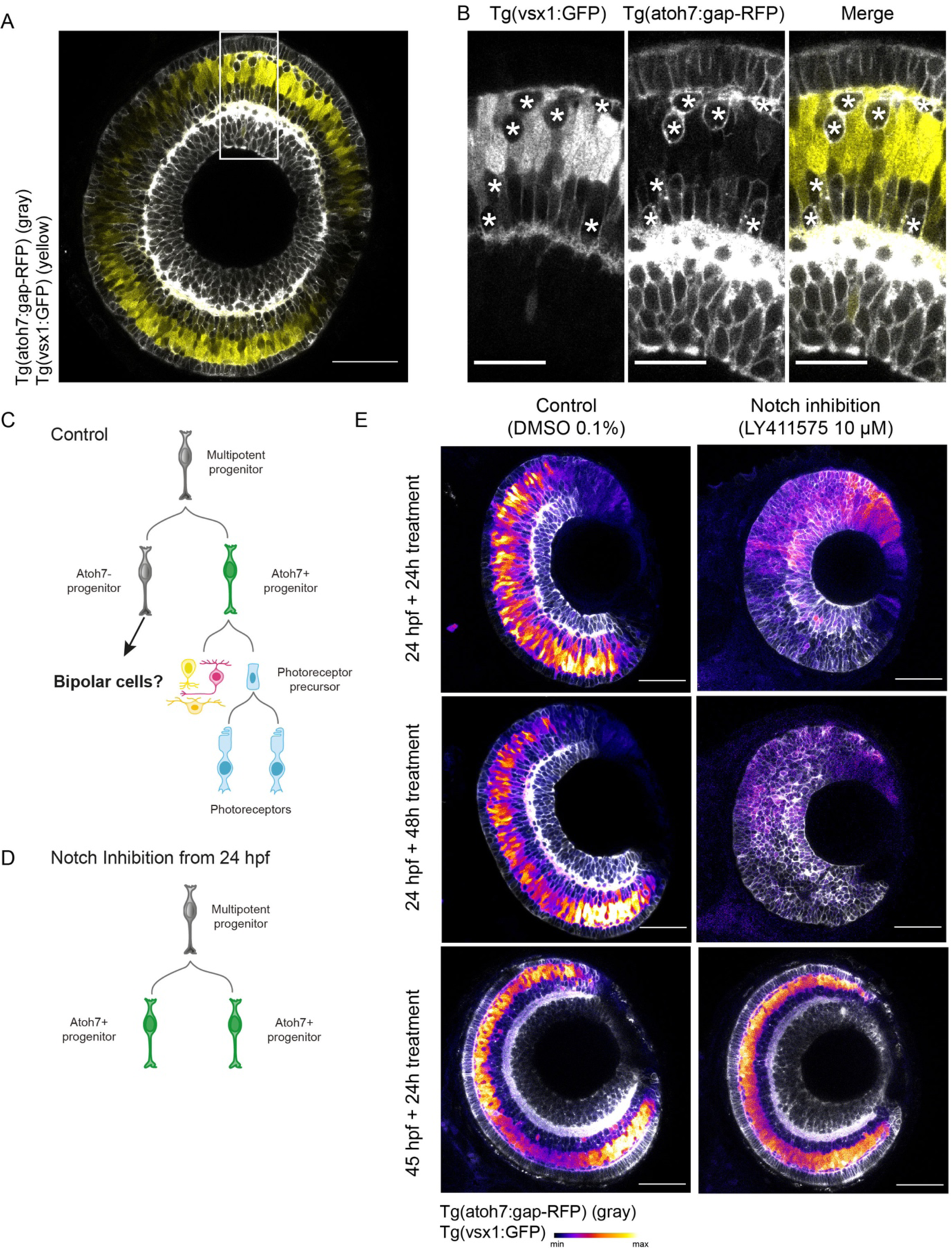
Bipolar cells originate from the Atoh7-negative sister cell of the Atoh7+ progenitor. A) Double transgenic line Tg(atoh7:gapRFP),Tg (vsx1:GFP) labelling Atoh7+ neurons (grey) and Bipolar cells (Vsx1+, yellow) at 60 hpf. Scale bar 50 µm. B) Close-up of (A) Tg (vsx1:GFP) (left), Tg(atoh7:gapRFP) (center) and their overlap (right). Stars indicate Atoh7+ Vsx1-cells. Scale bar 20 µm. C) Schematic of asymmetric division of multipotent progenitors that give rise to an Atoh7+ progenitor and an Atoh7-progenitor. D) Schematics of symmetric multipotent progenitor divisions that produce two Atoh7+ progenitors, occurring upon Notch inhibition from 24 hpf. E) Notch inhibition experiment. Left images show controls, right images show the Notch inhibition conditions with 10 µM LY411575 from 24 hpf and from 45 hpf. Treatment windows are indicated in the figure. Scale bar 50 µm.

**Supplementary Figure 3.**
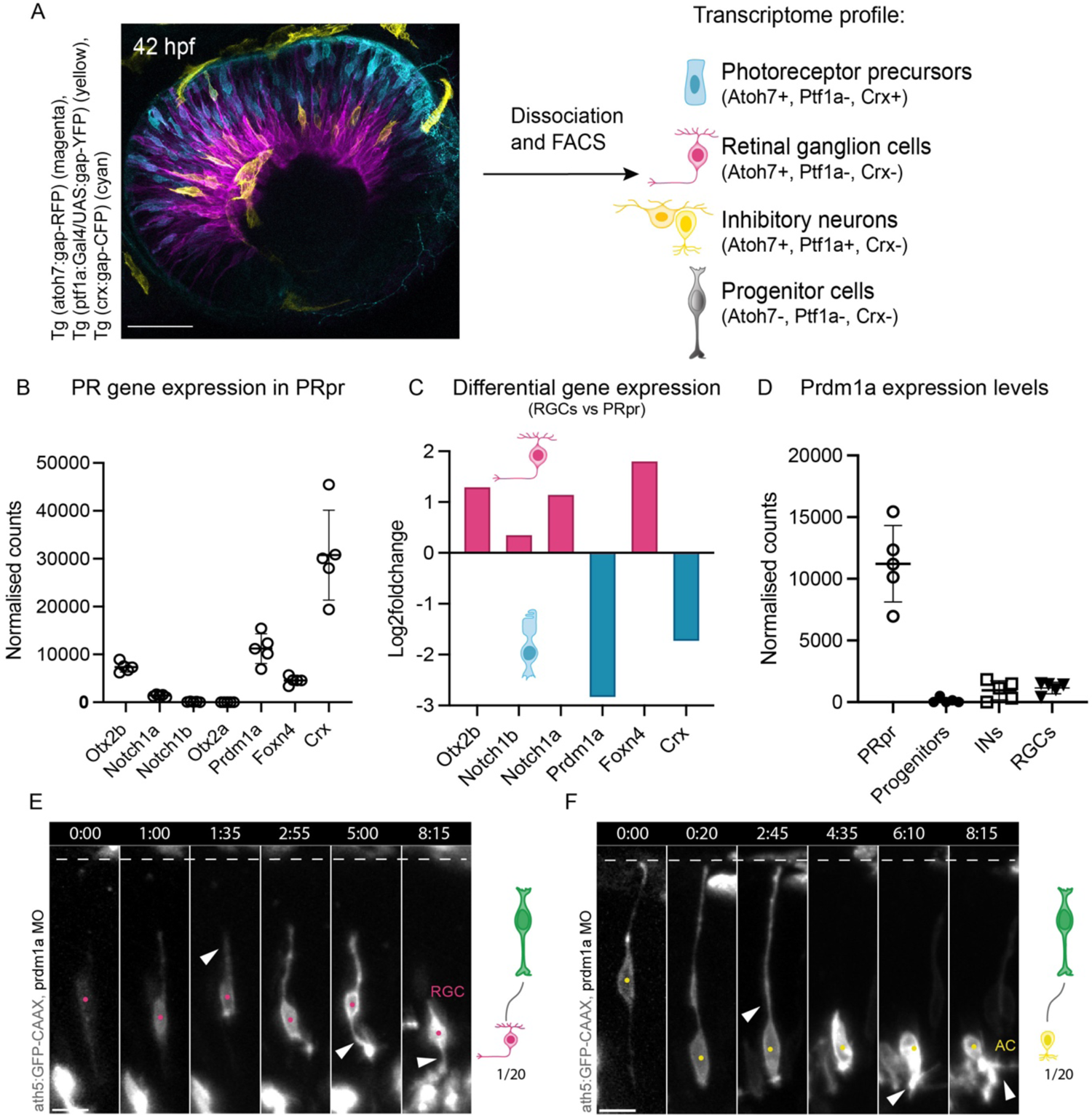
Prdm1a knockdown depletes PRs. A) Strategy for transcriptomics of PRpr, RGCs, INs as well as progenitor cells at 42 hpf. Atoh7+ cells (magenta), inhibitory neurons (yellow) and photoreceptors (cyan). Different cell types were selected based on the expression of different fluorophore combinations. B) Normalised counts of PR-related genes in the PRpr population from transcriptomics experiment. C) Differential expression of the same PR-related genes in the PR vs RGC comparison at 42 hpf. Magenta bar plots show genes enriched in the RGC population; cyan bar plots show genes enriched in the PR population. D) Normalised counts of Prdm1a expression levels in the PRpr, progenitors, INs and RGCs populations. E) Montage of Atoh7+ progenitor generating an RGC (magenta dot) without dividing. Dashed line indicates the apical side, arrowhead points at first retraction of the apical process (t = 1:35), then axon. ath5:GFP-CAAX (Atoh7, grey), scale bar 10 µm. F) Montage of Atoh7+ progenitor generating an AC (yellow dot) without dividing. Dashed line indicates the apical side, arrowhead points to first retraction of the apical process (t = 2:45), then to dendrites (t = 6.10 and t = 8.15). ath5:GFP-CAAX (Atoh7, grey), scale bar 10 µm.

**Supplementary figure 4.**
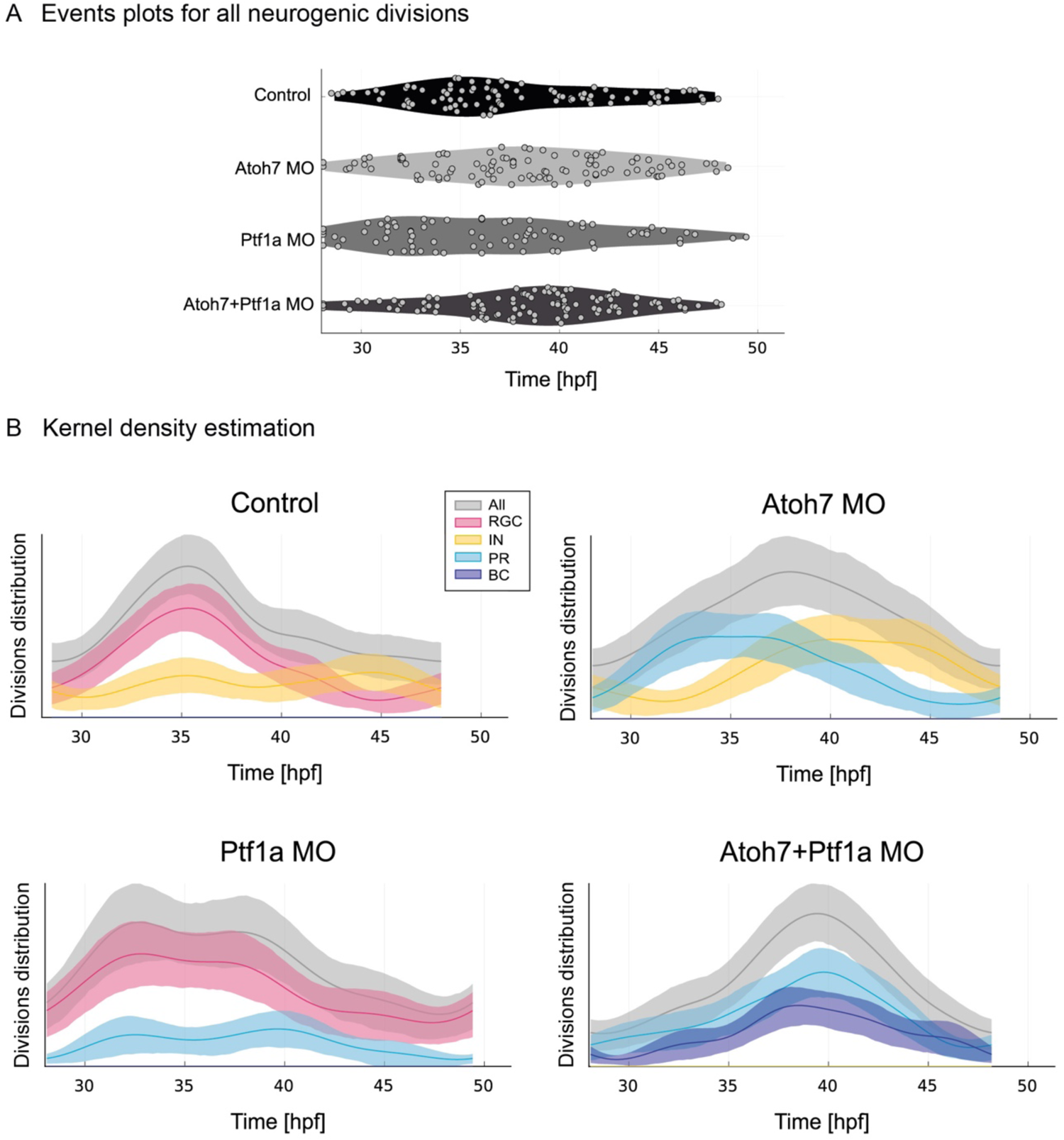
Absence of key TFs does not alter overall lineage topology. A) Distribution of each neurogenic Atoh7+ division from 28 hpf in control, Atoh7 morphants, Ptf1a morphants and double Atoh7/Ptf1a morphants. Time in hpf. Kolmogorov Smirnov test to compare distributions. Control vs Atoh7 morphant, p = 0.0836 (ns), Control vs Ptf1a morphant, p = 0.1369 (ns), Control vs Atoh7 + Ptf1a morphant, p = 0.0023. B) Kernel Density Estimation (KDE) of the distribution of events over time for each fate (coloured plots) and for the total distribution of Atoh7+ divisions (grey plots) during development in control, Atoh7 morphants, Ptf1a morphants and double Atoh7/Ptf1a morphants. Time in hpf. Mean (dark line) and 95% confidence interval (thick transparent stripe) are plotted.

## Video legends

**Video 1:** Division patterns of Atoh7+ retinal progenitors in controls.

*Part 1*: Asymmetric division of an Atoh7+ progenitor generating one RGC and one PRpr. Cells are labelled using ath5:GFP-CAAX (grey). Time is shown in minutes. Magenta and cyan dots label RGC and PRpr, respectively. Arrow points to RGC axon.

*Part 2*: Asymmetric division of an Atoh7+ progenitor generating one inhibitory neuron (HC) and one PRpr. Cells are labelled using ath5:GFP-CAAX (grey). Time is shown in minutes.

Orange and cyan dots label HC and PRpr, respectively.

*Part 3*: Asymmetric division of an Atoh7+ progenitor generating one inhibitory neuron (AC) and one PRpr. Cells are labelled using ath5:GFP-CAAX (grey). Time is shown in minutes.

Yellow and cyan dots label AC and PRpr, respectively. Arrow points to AC basal dendrites.

**Video 2:** Division patterns of Atoh7+ retinal progenitors in Prdm1a morphants.

Asymmetric division of an Atoh7+ progenitor generating one inhibitory neuron and one cell of undefined state. Cells are labelled using ath5:GFP-CAAX (grey). Time is shown in minutes. Orange and purple dots label Inhibitory neuron and cell of unknown state, respectively. Arrow points towards dynamic basal process of the cell of unknown state.

**Video 3:** Division patterns of Atoh7+ retinal progenitors in Atoh7/Ptf1a double morphants.

*Part 1*: Symmetric division of an Atoh7+ progenitor generating two PRpr. Cells are labelled using ath5:GFP-CAAX (grey). Time is shown in minutes. Cyan dots label the two PRprs.

*Part 2*: Asymmetric division of an Atoh7+ progenitor generating one bipolar cell (BC) and one PRpr. Cells are labelled using ath5:GFP-CAAX (grey). Time is shown in minutes. Blue and cyan dots label BC and PRpr, respectively.

## Author contributions

Conceptualization, E.N., T.B. and C.N.; Methodology, E.N., J.K., T.B., M.R.M., C.Z., and C.N.; Software, E.N and T.B.; Investigation, E.N., J.K., T.B., M.R.M.; Writing – Original Draft, E.N., T.B. and C.N.; Writing – Review & Editing, E.N., J.K., T.B., M.R.M., C.Z., and C.N.; Visualisation: E.N., T.B. and C.N.; Supervision, E.N., C.Z., and C.N; Funding Acquisition, C.Z., and C.N..

## Acknowledgements

We thank the Norden lab, the Zechner lab, the Modes lab and Pablo Sartori for fruitful discussions on the project. We are grateful to Anne Grapin-Botton and William A. Harris for helpful comments on the manuscript. Sylvia Kaufmann, Heike Hollak, Tânia Ferreira and João Coelho are thanked for technical help. We further thank the Computer Department, the Light Microscopy, Scientific Computing and Fish facilities of the Max Planck Institute of Molecular Cell Biology and Genetics as well as Advanced Imaging Unit and the Aquatic Facility at the Instituto Gulbenkian de Ciência for experimental support. We thank the Deep Sequencing Facility at the Genome Centre of the TUD for RNA-Seq and transcriptomic analysis.

